# The response of three-dimensional pancreatic alpha and beta cell co-cultures to oxidative stress

**DOI:** 10.1101/2021.09.07.459320

**Authors:** Mireille M.J.P.E. Sthijns, Timo Rademakers, Jolien Oosterveer, Thomas Geuens, Clemens A. van Blitterswijk, Vanessa L.S. LaPointe

## Abstract

The pancreatic islets of Langerhans have low endogenous antioxidant levels and are thus especially sensitive to oxidative stress, which is known to influence cell survival and behaviour. As bioengineered islets are gaining interest for therapeutic purposes, it is important to understand how their composition can be optimized to diminish oxidative stress. We investigated how the ratio of the two main islet cell types (alpha and beta cells) and their culture in three-dimensional aggregates could protect against oxidative stress. Monolayer and aggregate cultures were established by seeding the alphaTC1 (alpha) and INS1E (beta) cell lines in varying ratios, and hydrogen peroxide was applied to induce oxidative stress. Viability, oxidative stress, and the level of the antioxidant glutathione were measured. Both aggregation and an increasing prevalence of INS1E cells in the co-cultures conferred greater resistance to cell death induced by oxidative stress. Increasing the prevalence of INS1E cells also decreased the number of alphaTC1 cells experiencing oxidative stress in the monolayer culture. In 3D aggregates, culturing the alphaTC1 and INS1E cells in a ratio of 50:50 prevented oxidative stress in both cell types. Together, the results of this study lead to new insight into how modulating the composition and dimensionality of a co-culture can influence the oxidative stress levels experienced by the cells.

## 1. Introduction

Every cell has endogenous antioxidants that can protect against oxidants. Oxidants can be generated internally due to incomplete metabolism in mitochondria, for example as a result of decreased oxygen supply due to dissociation from the vasculature during tissue transplantation [1, 2]. In physiological situations, antioxidants can effectively protect against the damage induced by such oxidants [3–6]. However, when the endogenous level of antioxidants is too low or the oxidants overwhelm the antioxidants, this can lead to a disbalance between oxidants and antioxidants and induce oxidative stress. And while a low level of oxidants is necessary for a cell to function, a high level can lead to damage, diminished viability [7, 8], and can influence cell identity and fate [9]. For example, oxidants resulting from incomplete mitochondrial metabolism can influence stem cell fate decisions [9, 10]. For example, this is seen during the late stages of pancreatic development when a low level of oxygen inhibits differentiation towards the insulin-producing beta cells [11]. In general, alterations in the redox status have been shown to affect pancreatic islet function and modulate beta cell dysfunction [12–16] and viability [17].

Oxidative stress is important to consider in the pancreatic islets of Langerhans. These cells are highly sensitive to oxidative stress because they have very low endogenous antioxidant levels. Oxidative stress also appears to be an important factor in the progression towards type 1 diabetes [18]. And while the antioxidants thioredoxin, peroxiredoxin, and glutathione peroxidase 7 and 8 are known to enhance beta cell function, the sensitivity of alpha cells to oxidative stress and the effects of their interactions with beta cells remain unknown [19–23]. This becomes especially important as bioengineered islets, in which the composition of the different cell types can be modulated, are being considered for therapeutic applications [24, 25]. Furthermore, the role of the antioxidant glutathione (GSH) remains to be elucidated, and it has also been shown that oxidative stress sensitivity is different in a monolayer culture compared to a three-dimensional (3D) culture [26], which led us to wonder whether this would also be the case when alpha and beta cells are (co-)cultured in 3D aggregates.

We questioned how the ratio of the alpha and beta cells and their culture in monolayers compared to 3D aggregates would affect their susceptibility to oxidative stress. To investigate this, we made co-cultures of different ratios of alpha and beta cell lines; namely 100:0, 80:20, 50:50, 20:80 and 0:100 in both monolayer culture and 3D aggregates. We induced oxidative stress and investigated the effects on viability, the level of oxidative stress experienced by the different cell types, and the total level of the intracellular antioxidant GSH.

## 2. Materials and methods

### 2.1. Cell culture

A rat beta cell line (INS1E, AdexxBio C0018009) at passage 38–46 was cultured in RPMI (Gibco 21875) supplemented with 5% (v/v) fetal bovine serum (Sigma-Aldrich F7524), 1 mM sodium pyruvate, 10 mM HEPES, and 0.05 mM 2-mercaptoethanol. A mouse alpha cell line (alphaTC1 Clone 6; ATCC CRL-2934) at passage 54–62 was cultured in Dulbecco’s modified eagle medium (DMEM; Sigma-Aldrich D6046) supplemented with 10% (v/v) fetal bovine serum, 15 mM HEPES, 0.1 mM non-essential amino acids, 1.5 g/L sodium bicarbonate, and 2.0 g/L glucose. When the two cell lines were co-cultured, the respective media were combined in the same ratio as the cells. All cells were cultured in a humidified atmosphere containing 5% CO_2_ at 37°C, and were passaged 1–2 times a week with trypsinization. Medium was refreshed every two days.

### 2.2. Creation of alphaTC1-BFP2 and INS1E-mNeongreen

Mouse alpha cells were labelled with BFP2 and rat beta cells were labelled with mNeonGreen to distinguish the different cell types. To generate the vectors, the open reading frames of mTagBFP2 and mNeongreen2 were amplified by PCR from donor vectors pBAD-mTagBFP2 (Addgene #34632) and pSFFV_mNG2(11)1-10 (Addgene #82610), respectively, and ligated in a pLenti6.2 destination vector. After verification by Sanger sequencing, they were deposited as pLenti6.2_mTagBFP2 (Addgene #113725) and pLenti6.2_mNeonGreen2 (Addgene #113727). The plasmids were then co-transfected separately with third-generation lentiviral packaging and envelope vectors: pMD2.G (Addgene #12259), pRSV-Rev (Addgene #12253), and pMDLg/pRRE (Addgene #12251) into HEK293ft cells (Thermo Fisher R70007) using the PEIpro (VWR) transfection reagent. After 24 h, the viral supernatant was collected in either alphaTC1 or INS1E medium and filtered. Stable cell lines were generated by diluting the lentivirus 1:10 and 1:20 in cell type–specific medium and transducing the alphaTC1 and INS1E using pLenti6.2_mTagBFP2 and pLenti6.2_mNeonGreen2, respectively. Selection with 1 µg/ml puromycin dihydrochloride (Sigma-Aldrich) began 48 h after transduction and continued for at least 7–14 days. The transduction efficiency was assessed by flow cytometry and was determined to be 80.1% for INS1E cells and 89.7% for alphaTC1 cells.

### 2.3. Cell seeding

Three-dimensional aggregates were formed using low attachment microwell plates (STATARRAYS MCA96-16.224-PS-LA, 300MICRONS, Karlsruhe, Germany). To prepare the plates, an isopropanol series (70, 50, 30%) was used for deaeration, after which they were washed in PBS, coated overnight with 3% (w/v) Pluronics F-127 (Sigma-Aldrich), and washed again in PBS. The 3D aggregates were formed by seeding 169,000 cells per well in the following ratios of INS1E to alphaTC1: 0:100, 20:80, 50:50, 80:20 and 100:0. This resulted in approximately 169 aggregates per well of a size of 1000 cells each (which approximates the size and function of one physiological islet equivalent (1 IEQ) [27]). After seeding, the plate was centrifuged for 3 min at 100 × g. For the monolayer (co-)culture of INS1E and alphaTC1, the cells were seeded in the same ratios for a total of 400,000 cells per well in black 96-well tissue culture plates (Falcon). The 3D aggregates were cultured for 5 days and the monolayers were cultured for 24 h prior to starting experiments.

### 2.4. Induction of oxidative stress

In order to induce oxidative stress, both the monolayer cultures and 3D aggregates were exposed to 0, 20, 100, 500, 1000, or 2000 μM hydrogen peroxide (H_2_O_2_; Sigma-Aldrich) for 30 min in exposure medium. For the INS1E cells, the exposure medium comprised SILAC medium (Gibco, A2494201) supplemented with 1 mM sodium pyruvate, 10 mM HEPES, 3.0 g/L glucose, and 2 mM _L_-glutamine. For the alphaTC1 cells, the exposure medium comprised minimal essential DMEM (Gibco 11880) supplemented with 15 mM HEPES, 0.1 mM non-essential amino acids, 1.5 g/L sodium bicarbonate, and 3.0 g/L glucose. For measuring viability, the medium with H_2_O_2_ was replenished with fresh exposure medium before performing the assay. For measuring oxidative stress, the detection reagent was added to the medium containing H_2_O_2_.

### 2.5. Cell viability

The Cell Titer-Glo 2D and 3D viability assays (Promega) were used to measure viability by adding 100 μl of the Cell Titer-Glo reagent to an equal volume of exposure medium followed by an incubation for 5 min at 200 rpm on an orbital shaker. The signal was stabilized for 25 min and luminescence was measured on a microplate reader (CLARIOstar, BMG LABTECH) with a 0.25–1 second integration time.

### 2.6. Detection of oxidative stress

To detect cells experiencing oxidative stress, they were incubated with CellROX Deep Red Reagent (Thermo Fisher) in exposure medium for 30 min. Wheat germ agglutinin (WGA) Alexa Fluor 594 conjugate (Thermo Fisher) was added to the medium for the final 15 min to enhance segmentation by microscopy. The cells were fixed for 10–15 min in 4% formaldehyde in PBS at room temperature and were imaged within 24 h on a Nikon Ti widefield microscope equipped with a CrEST Optics spinning disk unit and a Lumencor SpectraX. Three-dimensional aggregates were imaged at 20× magnification (CFI Plan Apochromat λ) in confocal mode (pinhole size: 70 μm) using a piezo z-stage to sequentially image a full z-stack per channel. Z-stacks of 200 μm depth were taken with a step size of 8 μm. Images were acquired using the following channels: alphaTC1: Ex 395/25 nm, Em 447/60 nm; INS1E: Ex 470/24 nm, Em 520/35 nm; WGA: Ex 550/15 nm, Em 624/40 nm; CellROX: Ex 640/30 nm, Em 692/40 nm. The CellROX was imaged first to prevent potential light-induced activation of the probe.

### 2.7. Image analysis

The Nikon NIS Elements GA3 module (version 5.21) was used for automated image analysis and manual counting was performed if necessary. First, a background subtraction was performed in the channels for alphaTC1 and INS1E prior to 3D thresholding to enhance the detection of individual cells. After thresholding and skeletonization, the WGA signal was subtracted from the detected alphaTC1 and INS1E to improve segmentation. Mean fluorescence intensity of CellROX was measured within the remaining 3D volume of the individual alphaTC1 and INS1E cells. The percentage of the total number of cells and total alphaTC1 and INS1E cells that were positive for oxidative stress was reported. Since some alphaTC1 and INS1E were unlabelled (the transduction efficiency was approximately 80–90%), it should be noted that while the percentage of oxidative stress–positive cells is an accurate reflection of the whole population, the absolute number oxidative stress–positive cells is an underestimation.

### 2.8. GSH measurement

To determine the amount of GSH after induction of oxidative stress, the 3D aggregates were transferred to a microcentrifuge tube and centrifuged at 200 × g for 5 min. They were then washed in PBS, lysed in a 0.1 M potassium phosphate buffer (1.6:8.4 KH_2_PO_4_:K_2_HPO_4_; pH 7.5) supplemented with 10 mM EDTA disodium salt and 1% Triton-X, vortexed for 2 min, and incubated for 1 h at 4°C. Cells in monolayers were lysed in the same buffer for 30 min on ice, after which the cells were scraped and transferred to microcentrifuge tubes. All lysates were centrifuged at 14,000 × g for 10 min at 4°C. The total protein content of the supernatant was determined with a bicinchoninic acid assay (BCA; Pierce). To determine the amount of GSH, 300 μl of the supernatant was mixed 1:1 with 6% (w/v) sulfosalicylic acid (Sigma-Aldrich) in milliQ water. The enzymatic recycling method was applied to determine the amount of GSH present [28]. The absorbance was measured for 10 minutes at 37°C on a microplate reader (CLARIOstar, BMG LABTECH) at 412 nm and the slope of the curve was calculated. The slope (concentration of GSH over time) was corrected for the amount of protein measured.

### 2.9. Statistics

All data are presented as mean ± SEM. Statistical analyses were performed in GraphPad Prism (version 8.0). Comparisons were made using a two-way ANOVA with the Tukey test for multiple comparisons. Independent samples with equal variances were assessed for statistical significance with a t-test. P-values < 0.05 were considered statistically significant. N refers to the number of independent experiments and n to the number of replicates.

## 3. Results

### 3.1. 3D aggregation and increasing the percentage of INS1E cells conferred resistance to cell death induced by oxidative stress

INS1E and alphaTC1 cells were (co-)seeded in varying ratios (100:0, 80:20, 50:50, 20:80, and 0:100) in both monolayer cultures and 3D aggregates. As a first characterization of how the cells responded to oxidative stress, we exposed them to H_2_O_2_ and measured the resulting viability. We noted that both the dimensionality of the culture (*i.e.*, monolayer or 3D aggregate) and the ratio of INS1E to alphaTC1 affected viability after induced oxidative stress (Fig 1). The morphology of the cells in the monolayers or in the 3D aggregates was unaffected by the ratio of INS1E to alphaTC1 cells or the induction of oxidative stress when 0–1000 μM H_2_O_2_ was used (Fig S1, 2A and 4A).

**Figure 1:**
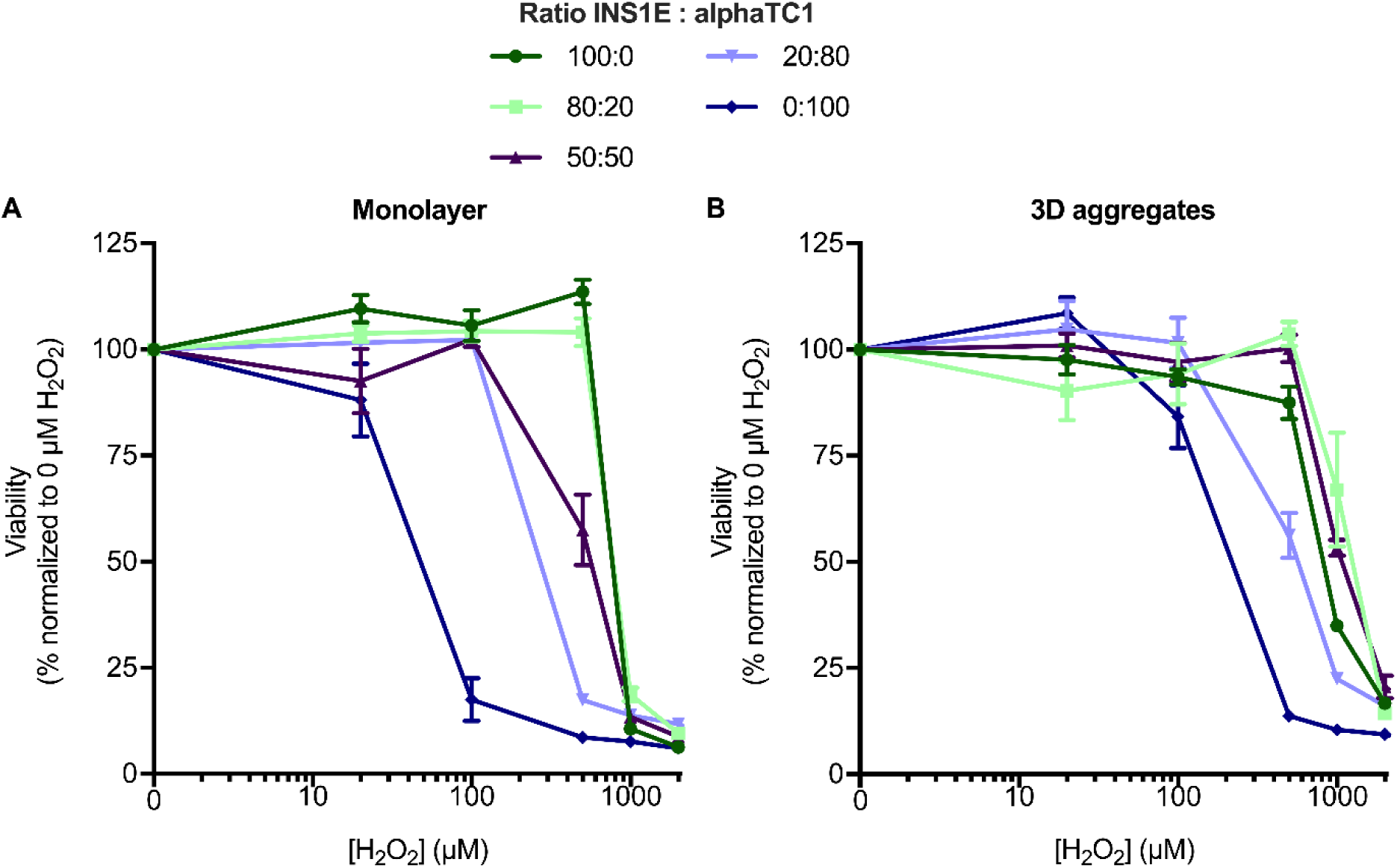
Cell viability in monolayer and 3D aggregate cultures after oxidative stress induced by H_2_O_2_. The percentage of viable cells is shown relative to the control that was not exposed to H_2_O_2_. A) In monolayer culture, the higher the percentage of INS1E cells included in the co-culture, the more H_2_O_2_ was necessary to induce a similar decrease in viability. B) In 3D aggregates of 80:20 and 50:50 INS1E:alphaTC1, the cells were more resilient to the loss of viability caused by H_2_O_2_ compared to the other ratios. N=3, A) n≥1; B) n≥2 and data are presented as mean ± SEM; all *p*-values are shown in Tables S1 and S2.

When comparing cells cultured in a monolayer (Fig 1A) or as 3D aggregates (Fig 1B), cells in the 3D aggregates were overall more resilient to reductions in viability caused by H_2_O_2_. For example, when INS1E and alphaTC1 cells were combined in a 50:50 ratio in a monolayer, it took 500 μM H_2_O_2_ to reduce their viability to 65% (*p*<0.001; Fig 1A, Table S1), but it took 1000 μM H_2_O_2_ for the same effect in 3D aggregates (*p*<0.001; Fig 1B, Table S1). We ruled out the possibility of these effects being due to limited diffusion in the 3D aggregate by staining and sectioning the aggregates to confirm the probe diffusion into their centres (Fig S2).

We then asked whether the ratio of INS1E to alphaTC1 cells would also affect the cells’ resilience to the H_2_O_2_-induced decreases we observed in cell viability. The outcome of this question was also influenced by whether the cells were cultured as a monolayer or in 3D aggregates.

In the monolayer (co-)culture, a higher percentage of INS1E cells generally meant there was less of a decrease in cell viability induced by oxidative stress (Fig 1A). For example, when alphaTC1 cells alone (without INS1E) were exposed to 100 μM H_2_O_2_, their viability was reduced to approximately 18% (*p*<0.001). Conversely, the same loss of viability in INS1E cells alone (without alphaTC1) was only observed after the induction of oxidative stress by a much higher (1000 μM) concentration of H_2_O_2_. For the other ratios of 80:20, 50:50 and 20:80 INS1E:alphaTC1, it was also seen that the higher the percentage of INS1E cells, the higher the concentration of H_2_O_2_ (1000 μM H_2_O_2_, 1000 μM H_2_O_2_, and 500 μM H_2_O_2_, respectively) was needed to be to induce a similar loss in viability to approximately 10% (all *p*<0.001).

In the 3D aggregate (co-)culture, the effect of combining INS1E with alphaTC1 was different. Overall, just like the cells in monolayers, having more INS1E cells in the aggregates provided resilience to the loss of viability induced by oxidative stress, but this trend did not directly relate to the percentage of INS1E (Fig 1B). For example, while increasing the ratio of INS1E from 0 to 20 to 100% increased their viability under the challenge of oxidative stress (Table S2, all *p*<0.001), the highest viability was observed when the percentage of INS1E in the 3D aggregate was either 50 or 80% (with no statistically significant difference between the two conditions).

Taken together, 3D aggregation and a higher prevalence of INS1E cells increased the resilience against a loss in viability induced by oxidative stress, but the ratio of the two cell types alone did not describe the resulting viability, especially in the 3D aggregates.

### 3.2. Increasing the prevalence of INS1E cells in monolayer culture diminished induced oxidative stress in alphaTC1 cells

Having seen that more INS1E cells in the monolayer co-culture protected against the overall loss of viability upon exposure to oxidative stress, we were interested to know whether this was due to the varying susceptibility of the two cell types to oxidative stress or whether the INS1E cells were conferring some protection to their neighbouring alphaTC1 cells. We hypothesized that the co-culture has a protective effect against the induction of oxidative stress.

To study this, we measured the level of intracellular oxidative stress when INS1E and alphaTC1 cells were seeded in different ratios (Fig S2 and Table S3 and S4). Regardless of their prevalence in the co-culture, approximately the same percentage of INS1E cells (from 42–60%; *p*>0.05) were positive for oxidative stress when it was induced by 500 μM H_2_O_2_ (Fig 2B; Table S5). This implies that the presence of alphaTC1 cells had no effect on the oxidative stress experienced by the INS1E cells.

**Figure 2:**
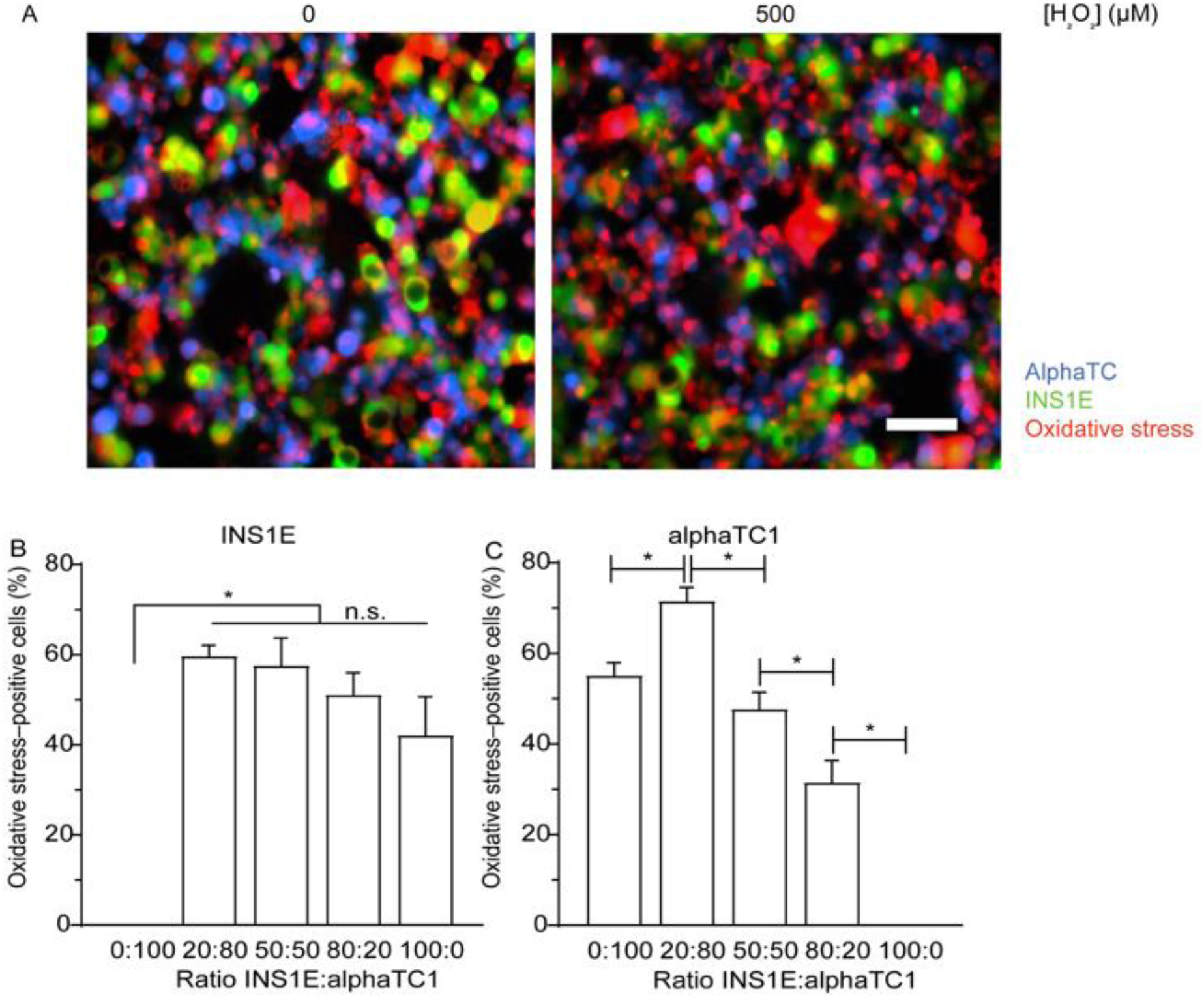
Oxidative stress in INS1E and alphaTC1 cells in monolayer culture. A) H_2_O_2_ in a concentration of 500 μM increased the number of oxidative stress positive (red) INS1E (green) and alphaTC1 (blue) cells. B) Varying the ratio of INS1E and alphaTC1 cells in monolayer culture exposed to 500 μM H_2_O_2_ did not change the percentage of oxidative stress–positive INS1E cells. C) Increasing the percentage of INS1E cells to 80% in a monolayer culture exposed to 500 μM H_2_O_2_ decreased the percentage of oxidative stress–positive alphaTC1 cells. N=3, n≥1 and data are presented as mean ± SEM; * *p*≤0.05; all p-values are shown in Tables S5 and S6.

Conversely, there was an effect on how the alphaTC1 cells experienced oxidative stress depending on how many INS1E cells were present in the monolayer co-culture. As a baseline, without INS1E cells in the co-culture, 55.1% of alphaTC1 cells were positive for oxidative stress when exposed to H_2_O_2_. When the culture comprised 20% INS1E cells, the percentage of alphaTC1 cells positive for oxidative stress increased to 71.6% (*p*<0.05; Fig 2C and Table S6). However, further increasing the prevalence of INS1E cells to 50% decreased the percentage of alphaTC1 cells that were positive for oxidative stress (47.7%; *p*<0.05). This trend continued when 80% of the culture comprised INS1E cells, which resulted in 31.5% of oxidative stress–positive alphaTC1 cells (*p*<0.05). A similar trend was seen at other H_2_O_2_ concentrations (from 20–1000 μM; Fig S3 and Table S5 and S6).

Overall, we conclude that in a monolayer co-culture, increasing the prevalence of INS1E cells can decrease the percentage of alphaTC1 cells that experience oxidative stress induced by H_2_O_2_, while increasing the prevalence of alphaTC1 cells did not have an effect on the percentage of oxidative stress–positive INS1E cells.

### 3.3. GSH decreased in oxidative stressed cells independent of the ratio of INS1E and alphaTC1 cells

In order to explain why increasing the prevalence of INS1E cells diminished the percentage of alphaTC1 cells experiencing oxidative stress in a monolayer co-culture but not the other way around, we sought a molecular basis of this observation. Therefore, we measured the intracellular level of GSH when alphaTC1 and INS1E cells were cultured alone or in combination at a ratio of 50:50.

We first measured the basal level of GSH (without the induction of oxidative stress by H_2_O_2_) because differing basal levels in the two cell types could explain why adding INS1E cells could diminish the oxidative stress experienced by the alphaTC1 cells, but not the other way around. However, there was no statistically significant difference in the intracellular level of GSH in cultures of alphaTC1 cells alone, INS1E cells alone, or a combination in the ratio 50:50 (Fig 3, Table S7).

**Figure 3:**
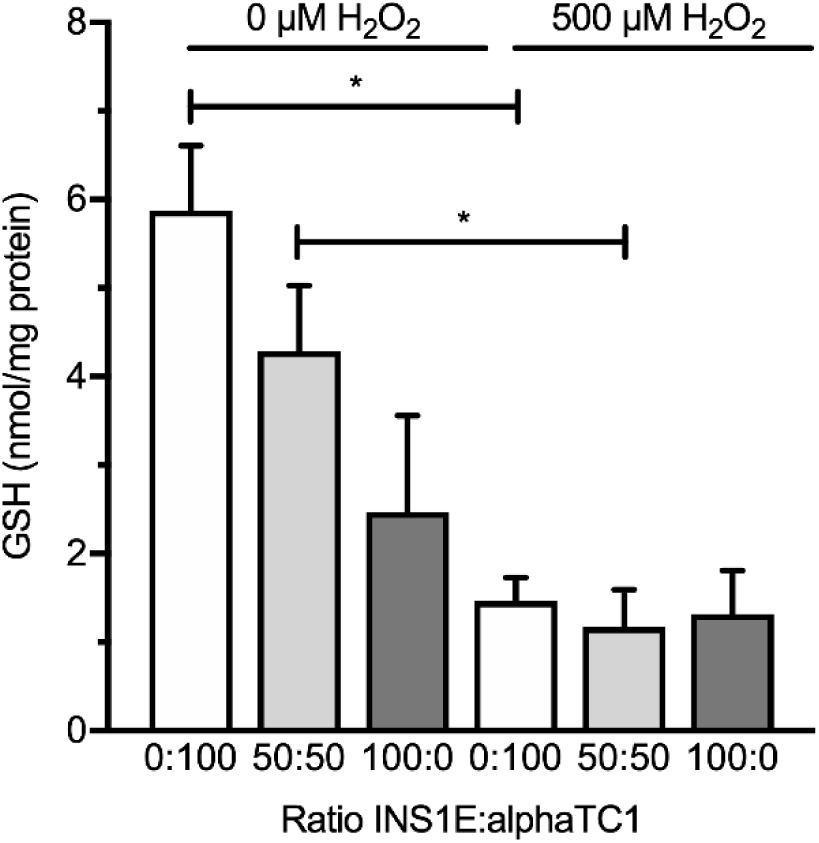
Intracellular GSH in monolayer cultures upon induced oxidative stress. Exposing the monolayer cultures of 0:100, 50:50 and 100:0 INS1E:alphaTC1 cells to 500 μM H_2_O_2_ only induced a significant decrease in intracellular GSH compared to a monolayer culture not exposed to H_2_O_2_ in the ratios of 50:50 and 0:100 INS1E:alphaTC1. N=3, n=2 and data are presented as mean ± SEM; * *p*≤0.05, all p-values are shown in Table S7.

We then measured how the basal level of GSH would be affected by the induction of oxidative stress with 500 μM H_2_O_2_. When INS1E cells were cultured alone, the induction of oxidative stress had no effect on the level of GSH (*p*=0.392 compared to the control of 0 μM H_2_O_2_). Conversely, when alphaTC1 cells were cultured alone or in combination with INS1E in a 50:50 ratio, the intracellular level of GSH was diminished upon induced oxidative stress (*p*<0.05 for both).

Together these experiments indicated that INS1E cells in a monolayer have such low levels of GSH that the addition of H_2_O_2_ did not further reduce them, whereas the GSH levels in alphaTC1 were reduced by the induction of oxidative stress.

### 3.4. Combining INS1E and alphaTC1 in 3D aggregates in a ratio of 50:50 prevented oxidative stress in both cell types

Having seen that, like in the monolayer culture, more INS1E cells in the aggregate co-cultures protected against the overall loss of viability upon exposure to oxidative stress, we went on to determine the varying susceptibility of the two cell types to oxidative stress. Observing that the prevalence of INS1E cells in the monolayer cultures was not directly related to the viability, we wondered how the cells may be influencing each other in aggregates.

Without alphaTC1 cells in the 3D aggregates, 38.4% of the INS1E cells experienced oxidative stress upon its induction by H_2_O_2_. With the addition of only 20% alphaTC1 cells, this percentage increased to 89.9% (Fig 4B; *p*<0.001), suggesting that the presence of alphaTC1 cells could affect the stress experienced by the INS1E cells. Interesting, when the co-culture was composed of 50% alphaTC1 cells, the number of oxidative stress–positive INS1E cells decreased again to 43.8% and did not differ significantly from the oxidative stress measured in 3D aggregates without alphaTC1 (*p*=0.828). Apparently, in 3D aggregates, combining INS1E and alphaTC1 in a ratio of 50:50 or culturing INS1E alone protects the INS1E cells against oxidative stress.

**Figure 4:**
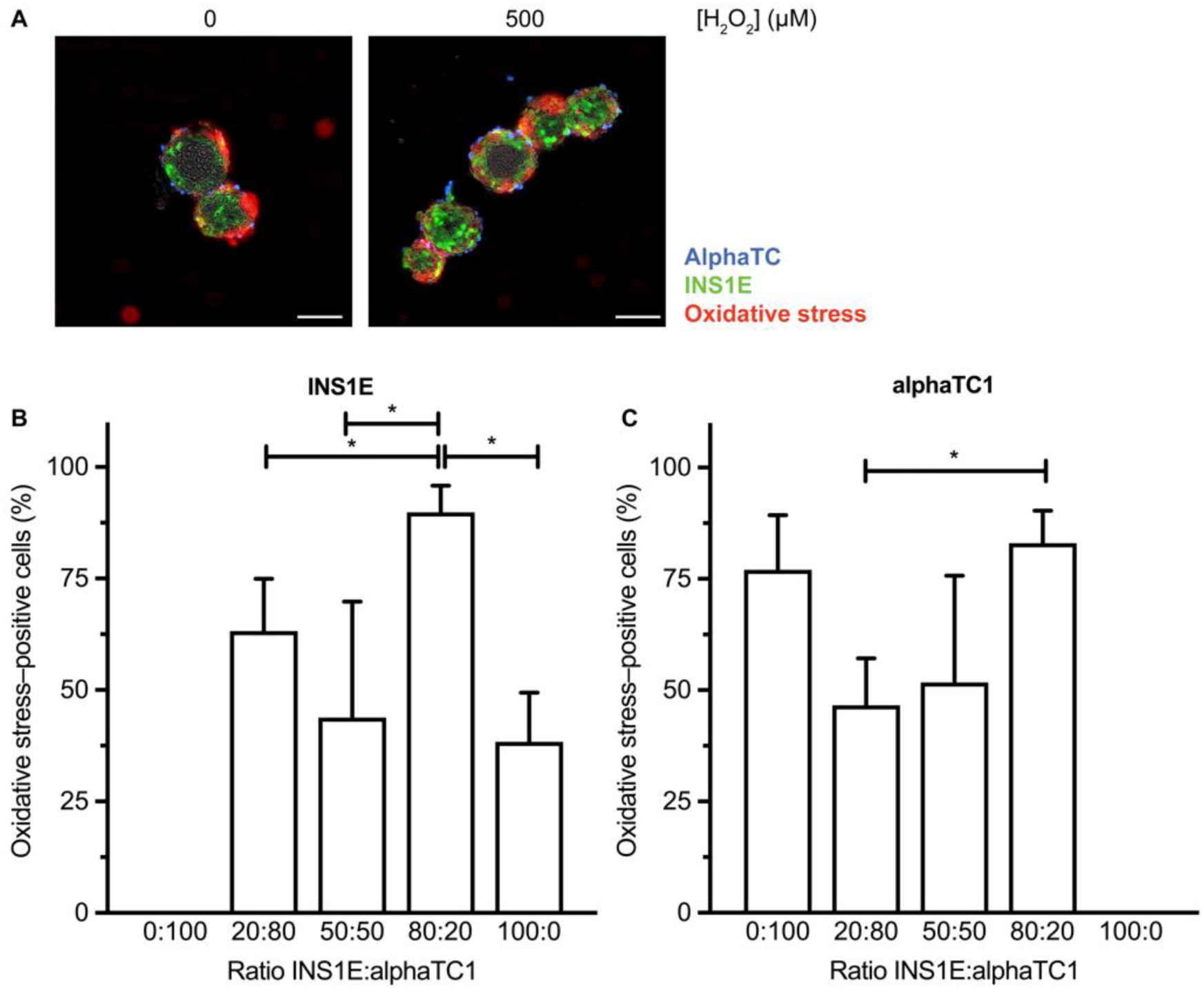
Oxidative stress in INS1E and alphaTC1 cells in 3D aggregates. A) Seeding INS1E and alphaTC1 cells in the ratios 100:0 and 50:50 in 3D aggregates exposed to 500 μM H_2_O_2_ decreased the percentage of oxidative stress–positive INS1E cells. B) Seeding INS1E and alphaTC1 in the ratio 20:80 in 3D aggregates exposed to 500 μM H_2_O_2_ decreased the percentage of oxidative stress–positive alphaTC1 cells compared to seeding 3D aggregates in the ratio 80:20 INS1E:alphaTC1. N=3, n≥2 and data are presented as mean ± SEM; * *p*≤0.05, all p-values are shown in Tables S8 and S9.

When the alphaTC1 cells were cultured alone, 77.0% of them experienced oxidative stress when exposed to 500 μM H_2_O_2_, a number significantly higher than what we measured in INS1E cells cultured alone (38.4%; *p*<0.05). Upon the addition of INS1E cells to the 3D aggregates, the percentage of alphaTC1 cells positive for oxidative stress decreased. For example, adding 20% of INS1E cells led to only 46.6% of alphaTC1 cells experiencing oxidative stress (Fig 4C). When the 3D aggregate contained 50% INS1E cells, only 51.7% of alphaTC1 cells experienced oxidative stress. This effect was reversed when the INS1E cells comprised 80% of the 3D aggregate, leading to 83.1% of alphaTC1 cells experiencing oxidative stress. Taken together, combining INS1E and alphaTC1 in a ratio of 50:50 protected both cell types against oxidative stress. A similar effect was seen at other H_2_O_2_ concentrations ranging from 20–100 μM (Fig S5 and Table S8 and 9).

Overall, we conclude that in 3D aggregates exposed to 500 μM H_2_O_2_, a 50:50 ratio decreased the percentage of both oxidative stress–positive alphaTC1 cells as well as INS1E cells.

### 3.5. 3D aggregates of INS1E had more GSH than aggregates of alphaTC1

To explain why combining INS1E and alphaTC1 cells in a ratio of 50:50 in 3D aggregates protected both cell types against oxidative stress, we measured the intracellular level of endogenous antioxidant GSH. In 3D aggregates of INS1E cells alone, we measured a higher GSH level than in aggregates of alphaTC1 cells alone (Fig 5; Table S10) (*p*<0.05). In addition, the intracellular level in a 3D aggregate in a ratio of 50:50 INS1E compared to alphaTC1 cells was higher (92.83 nmol/mg protein) than the GSH level in an aggregate of INS1E or alphaTC1 alone, although this was not statistically significant (Fig 5; Table S10). All in all, GSH could protect the 3D aggregates of INS1E cells alone and therefore explain why fewer oxidative stress–positive cells were measured compared to 3D aggregates of alphaTC1 cells alone when oxidative stress was induced (Fig 4).

**Figure 5:**
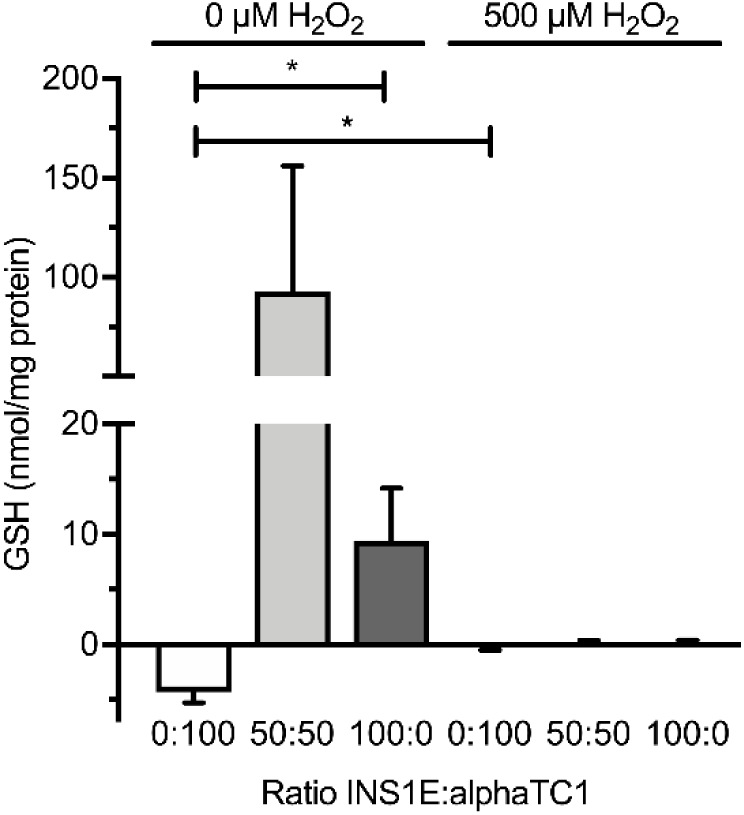
Intracellular GSH in 3D aggregates upon induced oxidative stress. A) The intracellular GSH in 3D aggregate of INS1E cells alone was significantly lower than in alphaTC1 cells alone. B) Exposing the 3D aggregates to 500 μM H_2_O_2_ only significantly affected the intracellular GSH in the ratio 0:100 INS1E:alphaTC1. N=3, n=2 and data are presented as mean ± SEM; * *p*≤0.05, all p-values are shown in Table S10.

## 4. Discussion

Every cell has a balance between oxidants and antioxidants, but in case of a disbalance, this can lead to oxidative stress that can induce cellular damage. Pancreatic islets are very sensitive to this damage because of their low intracellular antioxidant levels. To prevent this cellular damage, we questioned whether the ratio of the main islet cell types, alpha and beta cells, and culturing their interactions in 3D could protect against oxidative stress. To investigate this, we cultured the alpha and beta cell lines in different ratios of INS1E:alphaTC1 (100:0, 80:20, 50:50, 20:80 and 0:100), both in a monolayer and 3D aggregate (co-)cultures and exposed the cells to H_2_O_2_ to induce oxidative stress.

In summary, increasing the number of INS1E cells in a monolayer co-culture led to more resistance to decreases in viability induced by oxidative stress, whereas the viability was the highest in 3D aggregates comprising 50% and 80% INS1E cells (Fig 1). Increasing the prevalence of INS1E cells decreased the percentage of oxidative stress–positive alphaTC1 cells (Fig 2). In contrast, the percentage of oxidative stress–positive INS1E cells was not affected by the prevalence of alphaTC1 cells (Fig 2). Looking at the levels of intracellular GSH, the addition of H_2_O_2_ to a monolayer culture decreased the intracellular level of GSH in alphaTC1 cells alone and when they were in a 50:50 co-culture with INS1E cells (Fig 3). In 3D aggregates, a ratio of 50:50 INS1E:alphaTC1 was protective against oxidative stress in both cell types (Fig 4). When alphaTC1 cells were cultured alone in 3D aggregates, more oxidative stress–positive cells were induced by H_2_O_2_ than when INS1E cells were cultured alone (Fig 4). The 3D aggregates of INS1E cells alone had a higher level of intracellular GSH than 3D aggregates of alphaTC1 cells alone (Fig 5).

Our finding that H_2_O_2_ decreased the viability of INS1E is consistent with other studies [23, 29, 30], as is our result that 3D aggregation protected against an oxidative stress–induced decrease in viability [31, 32]. Overall, the oxidative stress the cells experienced did not correspond to their viability, which is not unexpected [33]. For example, we found that a monolayer culture with 100% alphaTC1 cells had the lowest viability compared to other ratios of the co-culture, even though it did not have the highest percentage of oxidative stress–positive cells.

In monolayer culture, the addition of H_2_O_2_ to a culture of 100% INS1E cells did not further decrease their basal intracellular antioxidant GSH levels (Fig 3), which is in agreement with the finding that endogenous antioxidant levels are very low in beta cells [19]. This is how a small addition of oxidants therefore could already create a redox disbalance that results in oxidative stress [34]. In contrast, the addition of 500 μM H_2_O_2_ to alphaTC1 cells significantly decreased the intracellular level of GSH. Apparently, the alphaTC1 cells have a different level of intracellular GSH. This finding that the endogenous antioxidant GSH levels were different between different cell types is in agreement with a previous report [35].

In 3D aggregate culture, a ratio of 50:50 alphaTC1 and INS1E cells prevented oxidative stress in both cell types (Fig 4). A possible explanation for this specific ratio is it increased the number of heterogenous cell–cell interactions. Due to this heterogenous interaction, one cell type could protect the other cell type against oxidative stress. For example glucagon-like peptide exendin-4 from alpha cells has been shown to decrease oxidative stress in beta cells [36]. In addition, alpha cells are necessary to set the glycaemic setpoint in beta cells [37]. This protective heterogenous cell–cell interaction has also been seen when mesenchymal stem cells were co-cultured with endocrine cells [38–40].

Our finding that INS1E cells experience less oxidative stress in 3D aggregates upon H_2_O_2_ exposure than alphaTC1 cells (Fig 4) could not be explained by the fact that the INS1E cells have higher GSH levels (Fig 5). The knowledge that beta cells enhance their function via connexion36-mediated interactions [41, 42] indicates that beta cells may be better in sharing these protective endogenous antioxidants amongst each other.

## 5. Conclusions

In conclusion, INS1E cells in a monolayer could protect alphaTC1 cells against oxidative stress. Establishing a 3D niche at the specific ratio of 50:50 INS1E:alphaTC1 cells was essential for protecting both cell types against oxidative stress. As a future outlook, it would be interesting to validate this finding in the context of beta cell replacement strategies, where the cells are known to experience high levels of oxidative stress [31, 32, 43–45]. Engineered islets could be transplanted in the ratio of 50:50 beta:alpha cells to protect them against the oxidative stress they experience during transplantation and from biomaterials used for encapsulation [46]. This approach becomes a reasonable strategy as the differentiation of pluripotent stem cells into pancreatic endocrine cells can create engineered islets resembling their *in vivo* counterparts [47].

## Author Contribution Statement

All authors have read and approved the final manuscript. M.M.J.P.E. Sthijns performed the experiments and designed the experimental outline. M.M.J.P.E. Sthijns performed imaging and analysis with the help of T. Rademakers. T. Geuens made the mNeongreen labelled INS1E cells and the BFP2 labelled alphaTC1 cells. J. Oosterveer performed the experiments from Fig 3 and 5. M.M.J.P.E. Sthijns and V.L.S. LaPointe wrote the manuscript with input from C.A. van Blitterswijk.

## Acknowledgements

The authors thank Fredrik Wieland for his help with the labelled cells.

## Data availability

The data that support the findings of this study will be made openly available in Dataverse upon acceptance.

## Supplementary Information

**Table S1:**
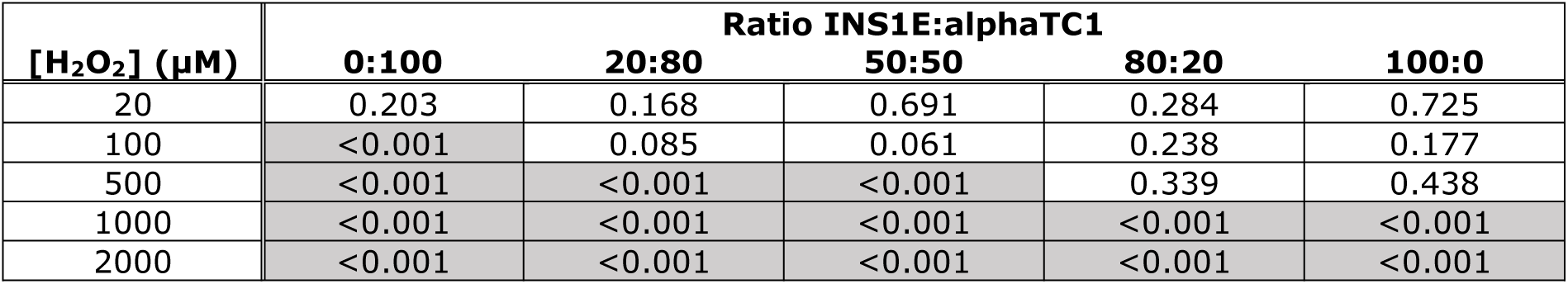
Statistical significance (t-test) of cell viability in monolayers upon induction of oxidative stress by H_2_O_2_ (20–2000 μM) compared to the control (0 μM H_2_O_2_) (from Fig 1A).

**Table S2:**
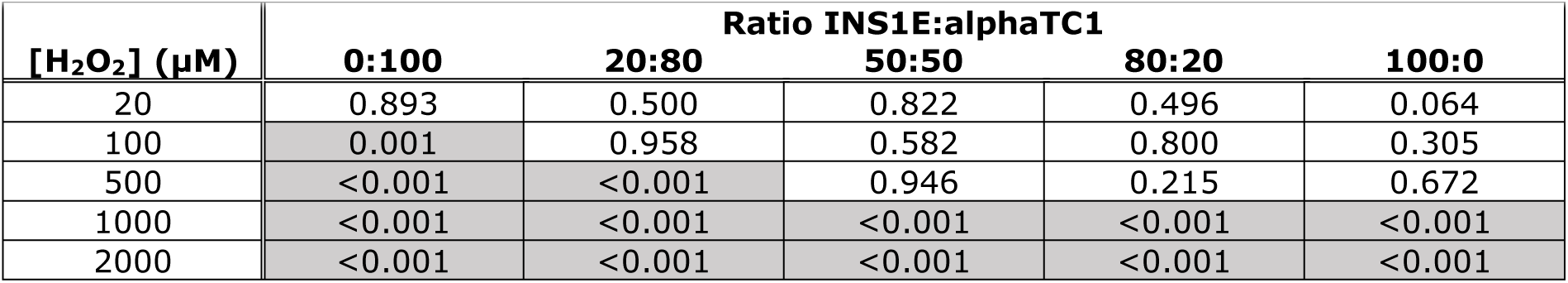
Statistical significance (t-test) of cell viability in 3D aggregates upon induction of oxidative stress by H_2_O_2_ (20–2000 μM) compared to the control (0 μM H_2_O_2_) (from Fig 1B).

**Table S3.** Statistical significance (t-test) of oxidative stress in monolayers upon induction by H_2_O_2_ (20–2000 μM) compared to the control (0 μM H_2_O_2_).

| [H<sub>2</sub>O<sub>2</sub>] (µM) | Ratio INS1E:alphaTC1 |  |  |  |  |
| --- | --- | --- | --- | --- | --- |
|  | 0:100 | 20:80 | 50:50 | 80:20 | 100:0 |
| 20 | 0.836 | 0.913 | 0.612 | 0.309 | 0.751 |
| 100 | 0.919 | 0.623 | 0.961 | 0.078 | 0.967 |
| 500 | 0.981 | 0.042 | 0.477 | 0.080 | 0.996 |
| 1000 | 0.656 | 0.006 | 0.201 | <0.001 | 0.324 |
| 2000 | 0.595 | 0.012 | 0.221 | <0.001 | 0.540 |

**Table S4.** Statistical significance (t-test) of oxidative stress in 3D aggregates upon induction by H_2_O_2_ (20–2000 μM) compared to the control (0 μM H_2_O_2_).

| [H<sub>2</sub>O<sub>2</sub>] (µM) | Ratio INS1E:alphaTC1 |  |  |  |  |
| --- | --- | --- | --- | --- | --- |
|  | 0:100 | 20:80 | 50:50 | 80:20 | 100:0 |
| 20 | 0.888 | 0.569 | 0.721 | 0.562 | 0.564 |
| 100 | 0.259 | 0.605 | 0.511 | 0.758 | 0.994 |
| 500 | 0.555 | 0.653 | 0.483 | 0.582 | 0.975 |
| 1000 | 0.516 | 0.624 | 0.290 | 0.683 | 0.379 |
| 2000 | 0.804 | 0.412 | 0.178 | 0.998 | 0.104 |

**Table S5.** Statistical significance (t-test) of the oxidative stress positive INS1E cells in monolayer co-cultures when exposed to 0–2000 μM H_2_O_2_.

|  |  | Ratio INS1E:alphaTC1 |  |  |  |  |
| --- | --- | --- | --- | --- | --- | --- |
|  |  | 0:100 | 20:80 | 50:50 | 80:20 | 100:0 |
| 0 $\mu\text{M}$ | 0:100 | -- | <0.001 | <0.001 | <0.001 | <0.001 |
|  | 20:80 | -- | -- | 0.045 | 0.045 | 0.864 |
|  | 50:50 | -- | -- | -- | 0.005 | 0.199 |
|  | 80:20 | -- | -- | -- | -- | 0.295 |
|  | 100:0 | -- | -- | -- | -- | -- |
|  |  | 0:100 | 20:80 | 50:50 | 80:20 | 100:0 |
| 20 $\mu\text{M}$ | 0:100 | -- | <0.001 | <0.001 | <0.001 | <0.001 |
|  | 20:80 | -- | -- | 0.144 | 0.385 | 0.734 |
|  | 50:50 | -- | -- | -- | 0.054 | 0.184 |
|  | 80:20 | -- | -- | -- | -- | 0.738 |
|  | 100:0 | -- | -- | -- | -- | -- |
|  |  | 0:100 | 20:80 | 50:50 | 80:20 | 100:0 |
| 100 $\mu\text{M}$ | 0:100 | -- | <0.001 | <0.001 | <0.001 | <0.001 |
|  | 20:80 | -- | -- | 0.178 | 0.547 | 0.459 |
|  | 50:50 | -- | -- | -- | 0.614 | 0.106 |
|  | 80:20 | -- | -- | -- | -- | 0.350 |
|  | 100:0 | -- | -- | -- | -- | -- |
|  |  | 0:100 | 20:80 | 50:50 | 80:20 | 100:0 |
| 500 $\mu\text{M}$ | 0:100 | -- | <0.001 | <0.001 | <0.001 | <0.001 |
|  | 20:80 | -- | -- | 0.701 | 0.114 | 0.147 |
|  | 50:50 | -- | -- | -- | 0.424 | 0.359 |
|  | 80:20 | -- | -- | -- | -- | 0.359 |
|  | 100:0 | -- | -- | -- | -- | -- |
|  |  | 0:100 | 20:80 | 50:50 | 80:20 | 100:0 |
| 1000 $\mu\text{M}$ | 0:100 | -- | <0.001 | <0.001 | <0.001 | <0.001 |
|  | 20:80 | -- | -- | 0.936 | <0.001 | <0.001 |
|  | 50:50 | -- | -- | -- | 0.030 | 0.001 |
|  | 80:20 | -- | -- | -- | -- | <0.001 |
|  | 100:0 | -- | -- | -- | -- | -- |
|  |  | 0:100 | 20:80 | 50:50 | 80:20 | 100:0 |
| 2000 $\mu\text{M}$ | 0:100 | -- | <0.001 | <0.001 | <0.001 | <0.001 |
|  | 20:80 | -- | -- | 0.714 | 0.002 | <0.001 |
|  | 50:50 | -- | -- | -- | 0.103 | 0.005 |
|  | 80:20 | -- | -- | -- | -- | <0.001 |
|  | 100:0 | -- | -- | -- | -- | -- |

**Supplementary Table 6.** Statistical significance (t-test) of the oxidative stress positive alphaTC1 cells in monolayer co-cultures when exposed to 0–2000 μM H_2_O_2_.

|  |  | Ratio INS1E:alphaTC1 |  |  |  |  |
| --- | --- | --- | --- | --- | --- | --- |
|  |  | 0:100 | 20:80 | 50:50 | 80:20 | 100:0 |
| 0 µM | 0:100 | -- | 0.217 | 0.012 | <0.001 | <0.001 |
|  | 20:80 | -- | -- | <0.001 | <0.001 | <0.001 |
|  | 50:50 | -- | -- | -- | 0.097 | <0.001 |
|  | 80:20 | -- | -- | -- | -- | <0.001 |
|  | 100:0 | -- | -- | -- | -- | -- |
|  |  | 0:100 | 20:80 | 50:50 | 80:20 | 100:0 |
| 20 µM | 0:100 | -- | 0.264 | 0.115 | <0.001 | <0.001 |
|  | 20:80 | -- | -- | 0.006 | <0.001 | <0.001 |
|  | 50:50 | -- | -- | -- | 0.009 | <0.001 |
|  | 80:20 | -- | -- | -- | -- | <0.001 |
|  | 100:0 | -- | -- | -- | -- | -- |
|  |  | 0:100 | 20:80 | 50:50 | 80:20 | 100:0 |
| 100 µM | 0:100 | -- | 0.174 | 0.037 | 0.015 | <0.001 |
|  | 20:80 | -- | -- | <0.001 | <0.001 | <0.001 |
|  | 50:50 | -- | -- | -- | 0.408 | <0.001 |
|  | 80:20 | -- | -- | -- | -- | <0.001 |
|  | 100:0 | -- | -- | -- | -- | -- |
|  |  | 0:100 | 20:80 | 50:50 | 80:20 | 100:0 |
| 500 µM | 0:100 | -- | 0.005 | 0.163 | 0.009 | <0.001 |
|  | 20:80 | -- | -- | <0.001 | <0.001 | <0.001 |
|  | 50:50 | -- | -- | -- | 0.029 | <0.001 |
|  | 80:20 | -- | -- | -- | -- | <0.001 |
|  | 100:0 | -- | -- | -- | -- | -- |
|  |  | 0:100 | 20:80 | 50:50 | 80:20 | 100:0 |
| 1000 µM | 0:100 | -- | 0.002 | 0.477 | 0.852 | <0.001 |
|  | 20:80 | -- | -- | <0.001 | 0.002 | <0.001 |
|  | 50:50 | -- | -- | -- | 0.367 | <0.001 |
|  | 80:20 | -- | -- | -- | -- | <0.001 |
|  | 100:0 | -- | -- | -- | -- | -- |
|  |  | 0:100 | 20:80 | 50:50 | 80:20 | 100:0 |
| 2000 µM | 0:100 | -- | 0.003 | 0.408 | 0.473 | <0.001 |
|  | 20:80 | -- | -- | 0.001 | 0.014 | <0.001 |
|  | 50:50 | -- | -- | -- | 0.134 | <0.001 |
|  | 80:20 | -- | -- | -- | -- | <0.001 |
|  | 100:0 | -- | -- | -- | -- | -- |

**Supplementary Table 7.** Statistical significance (t-test) of the intracellular GSH levels in monolayer co-cultures when exposed to 500 μM H_2_O_2_ (Fig 3).

| INS1E:alphaTC1 | | 0 $\mu\text{M}$ $\text{H}_2\text{O}_2$ | | | 500 $\mu\text{M}$ $\text{H}_2\text{O}_2$ | | |
| --- | --- | --- | --- | --- | --- | --- | --- |
|  |  | 0:100 | 50:50 | 100:0 | 0:100 | 50:50 | 100:0 |
| 0 $\mu\text{M}$ $\text{H}_2\text{O}_2$ | 0:100 | -- | 0.203 | 0.611 | 0.005 | -- | -- |
|  | 50:50 | -- | -- | 0.240 | -- | 0.022 | -- |
|  | 100:0 | -- | -- | -- | -- | -- | 0.392 |
| 500 $\mu\text{M}$ $\text{H}_2\text{O}_2$ | 0:100 | -- | -- | -- | -- | 0.832 | 0.807 |
|  | 50:50 | -- | -- | -- | -- | -- | 0.592 |
|  | 100:0 | -- | -- | -- | -- | -- | -- |

**Supplementary Table 8.** Statistical significance (t-test) of the oxidative stress positive INS1E cells in 3D aggregate co-cultures when exposed to 0–2000 μM H_2_O_2_.

|  |  | Ratio INS1E:alphaTC1 |  |  |  |  |
| --- | --- | --- | --- | --- | --- | --- |
|  |  | 0:100 | 20:80 | 50:50 | 80:20 | 100:0 |
| 0 $\mu\text{M}$ | 0:100 | -- | <0.001 | 0.009 | <0.001 | <0.001 |
|  | 20:80 | -- | -- | 0.023 | 0.234 | 0.037 |
|  | 50:50 | -- | -- | -- | <0.001 | 0.211 |
|  | 80:20 | -- | -- | -- | -- | <0.001 |
|  | 100:0 | -- | -- | -- | -- | -- |
|  |  | 0:100 | 20:80 | 50:50 | 80:20 | 100:0 |
| 20 $\mu\text{M}$ | 0:100 | -- | <0.001 | 0.002 | <0.001 | 0.005 |
|  | 20:80 | -- | -- | 0.206 | 0.020 | 0.042 |
|  | 50:50 | -- | -- | -- | <0.001 | 0.916 |
|  | 80:20 | -- | -- | -- | -- | <0.001 |
|  | 100:0 | -- | -- | -- | -- | -- |
|  |  | 0:100 | 20:80 | 50:50 | 80:20 | 100:0 |
| 100 $\mu\text{M}$ | 0:100 | -- | 0.005 | 0.021 | <0.001 | 0.004 |
|  | 20:80 | -- | -- | 0.849 | 0.011 | 0.466 |
|  | 50:50 | -- | -- | -- | 0.043 | 0.762 |
|  | 80:20 | -- | -- | -- | -- | <0.001 |
|  | 100:0 | -- | -- | -- | -- | -- |
|  |  | 0:100 | 20:80 | 50:50 | 80:20 | 100:0 |
| 500 $\mu\text{M}$ | 0:100 | -- | <0.001 | 0.009 | <0.001 | 0.004 |
|  | 20:80 | -- | -- | 0.463 | 0.033 | 0.141 |
|  | 50:50 | -- | -- | -- | 0.017 | 0.828 |
|  | 80:20 | -- | -- | -- | -- | <0.001 |
|  | 100:0 | -- | -- | -- | -- | -- |
|  |  | 0:100 | 20:80 | 50:50 | 80:20 | 100:0 |
| 1000 $\mu\text{M}$ | 0:100 | -- | <0.001 | 0.004 | <0.001 | <0.001 |
|  | 20:80 | -- | -- | 0.809 | 0.104 | 0.296 |
|  | 50:50 | -- | -- | -- | 0.156 | 0.655 |
|  | 80:20 | -- | -- | -- | -- | 0.003 |
|  | 100:0 | -- | -- | -- | -- | -- |
|  |  | 0:100 | 20:80 | 50:50 | 80:20 | 100:0 |
| 2000 $\mu\text{M}$ | 0:100 | -- | <0.001 | <0.001 | <0.001 | <0.001 |
|  | 20:80 | -- | -- | 0.660 | 0.157 | 0.478 |
|  | 50:50 | -- | -- | -- | 0.161 | 0.995 |
|  | 80:20 | -- | -- | -- | -- | 0.032 |
|  | 100:0 | -- | -- | -- | -- | -- |

**Supplementary Table 9.** Statistical significance (t-test) of the oxidative stress positive alphaTC1 cells in 3D aggregate co-cultures when exposed to 0–2000 μM H_2_O_2_.

|  |  | Ratio INS1E:alphaTC1 |  |  |  |  |
| --- | --- | --- | --- | --- | --- | --- |
|  |  | 0:100 | 20:80 | 50:50 | 80:20 | 100:0 |
| 0 $\mu\text{M}$ | 0:100 | -- | 0.609 | 0.046 | 0.704 | <0.001 |
|  | 20:80 | -- | -- | 0.120 | 0.377 | <0.001 |
|  | 50:50 | -- | -- | -- | 0.038 | 0.006 |
|  | 80:20 | -- | -- | -- | -- | <0.001 |
|  | 100:0 | -- | -- | -- | -- | -- |
| 20 $\mu\text{M}$ | 0:100 | -- | 0.469 | 0.270 | 0.264 | <0.001 |
|  | 20:80 | -- | -- | 0.557 | 0.038 | <0.001 |
|  | 50:50 | -- | -- | -- | 0.053 | 0.002 |
|  | 80:20 | -- | -- | -- | -- | <0.001 |
|  | 100:0 | -- | -- | -- | -- | -- |
| 100 $\mu\text{M}$ | 0:100 | -- | 0.830 | 0.918 | 0.036 | 0.002 |
|  | 20:80 | -- | -- | 0.801 | 0.013 | 0.004 |
|  | 50:50 | -- | -- | -- | 0.186 | 0.004 |
|  | 80:20 | -- | -- | -- | -- | <0.001 |
|  | 100:0 | -- | -- | -- | -- | -- |
| 500 $\mu\text{M}$ | 0:100 | -- | 0.075 | 0.338 | 0.658 | <0.001 |
|  | 20:80 | -- | -- | 0.827 | 0.007 | <0.001 |
|  | 50:50 | -- | -- | -- | 0.134 | 0.001 |
|  | 80:20 | -- | -- | -- | -- | <0.001 |
|  | 100:0 | -- | -- | -- | -- | -- |
| 1000 $\mu\text{M}$ | 0:100 | -- | 0.407 | 0.659 | 0.724 | <0.001 |
|  | 20:80 | -- | -- | 0.927 | 0.133 | <0.001 |
|  | 50:50 | -- | -- | -- | 0.385 | 0.002 |
|  | 80:20 | -- | -- | -- | -- | <0.001 |
|  | 100:0 | -- | -- | -- | -- | -- |
| 2000 $\mu\text{M}$ | 0:100 | -- | 0.822 | 0.715 | 0.363 | 0.001 |
|  | 20:80 | -- | -- | 0.731 | 0.390 | <0.001 |
|  | 50:50 | -- | -- | -- | 0.862 | <0.001 |
|  | 80:20 | -- | -- | -- | -- | <0.001 |
|  | 100:0 | -- | -- | -- | -- | -- |

**Supplementary Table 10.** Statistical significance (t-test) of the intracellular GSH levels in 3D aggregate co-cultures when exposed to 500 μM H_2_O_2_ (Fig 5).

| INS1E:alphaTC1 | | 0 $\mu\text{M}$ $\text{H}_2\text{O}_2$ | | | 500 $\mu\text{M}$ $\text{H}_2\text{O}_2$ | | |
| --- | --- | --- | --- | --- | --- | --- | --- |
|  |  | 0:100 | 50:50 | 100:0 | 0:100 | 50:50 | 100:0 |
| 0 $\mu\text{M}$ $\text{H}_2\text{O}_2$ | 0:100 | -- | 0.199 | 0.048 | 0.016 | -- | -- |
|  | 50:50 | -- | -- | 0.258 | -- | 0.216 | -- |
|  | 100:0 | -- | -- | -- | -- | -- | 0.121 |
| 500 $\mu\text{M}$ $\text{H}_2\text{O}_2$ | 0:100 | -- | -- | -- | -- | 0.534 | 0.626 |
|  | 50:50 | -- | -- | -- | -- | -- | 0.951 |
|  | 100:0 | -- | -- | -- | -- | -- | -- |

**Figure S1:**
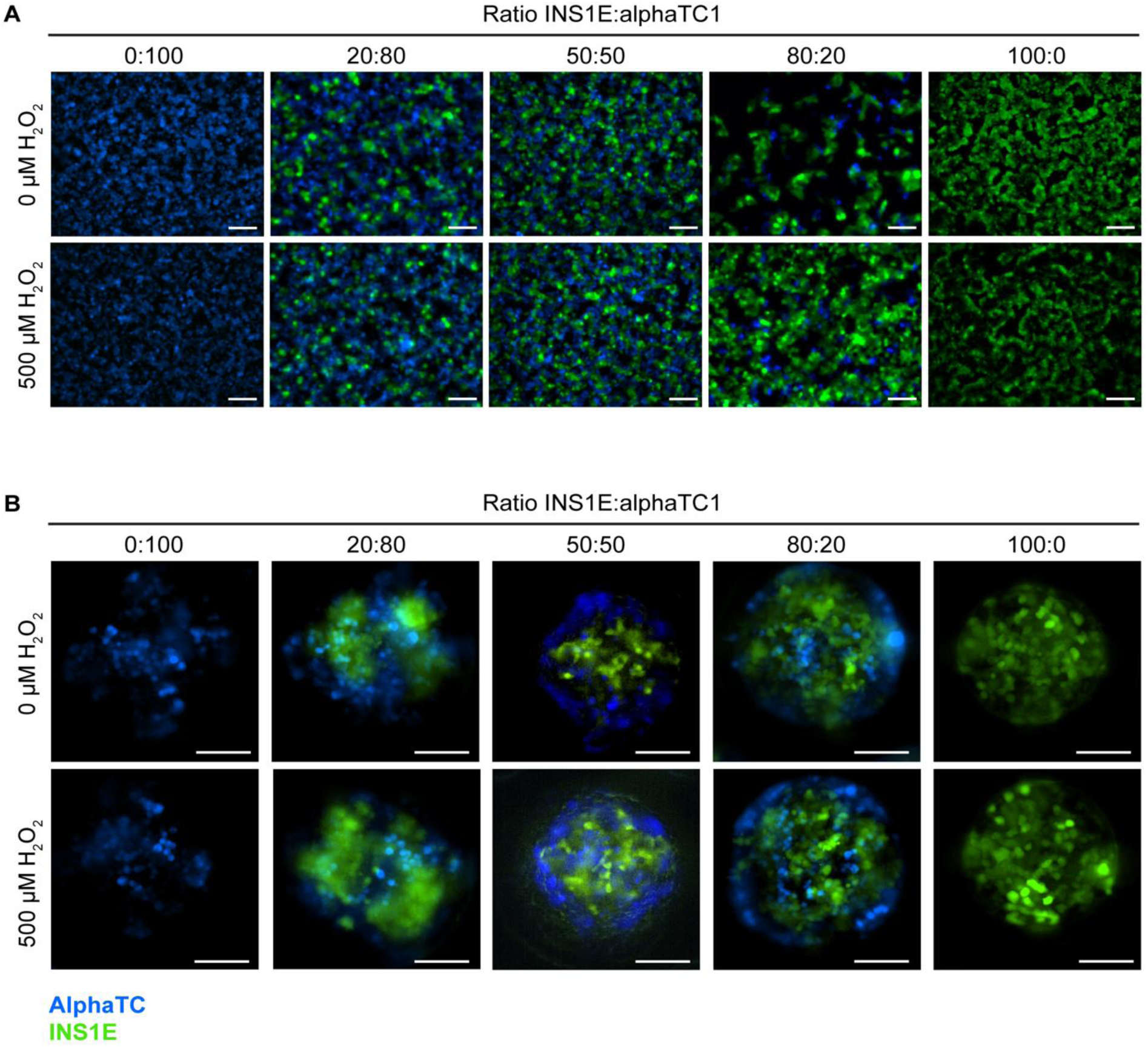
Morphology of cells in monolayer cultures and 3D aggregates. A) H_2_O_2_ did not affect the morphology of the INS1E and alphaTC1 cells in the different ratios in a monolayer. B) H_2_O_2_ also did not affect the morphology of the INS1E and alphaTC1 cells in the different ratios of the 3D aggregates. N=3 and a typical example is shown.

**Figure S2:**
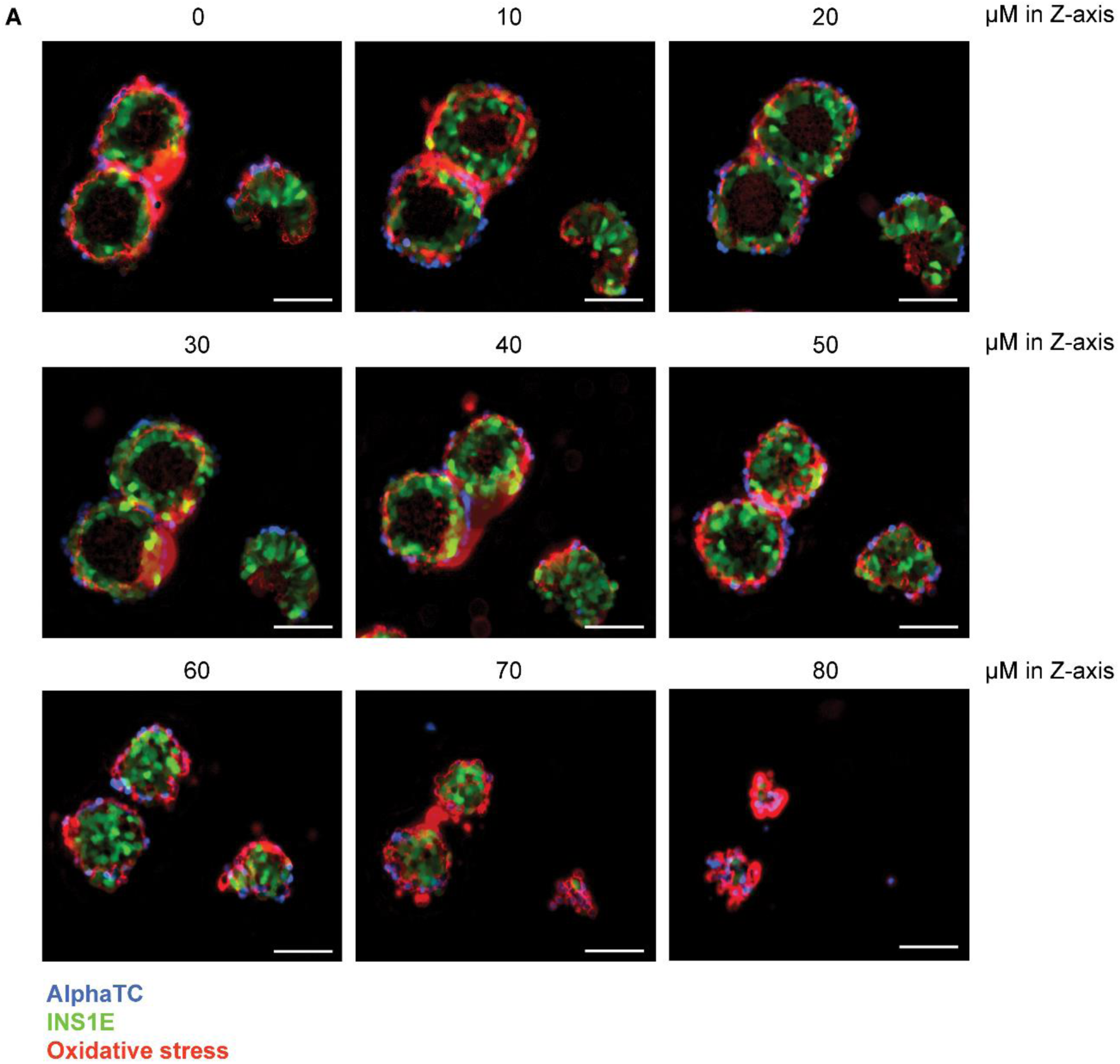
Sections of 3D aggregates. A) Increasing H_2_O_2_ increased the oxidative stress–positive cells (red). B) When exposed to 1000 μM H_2_O_2_, independent of the Z-stack number, cells within the 3D aggregate were coloured red because of the oxidative stress induced. A typical example is shown.

**Figure S3:**
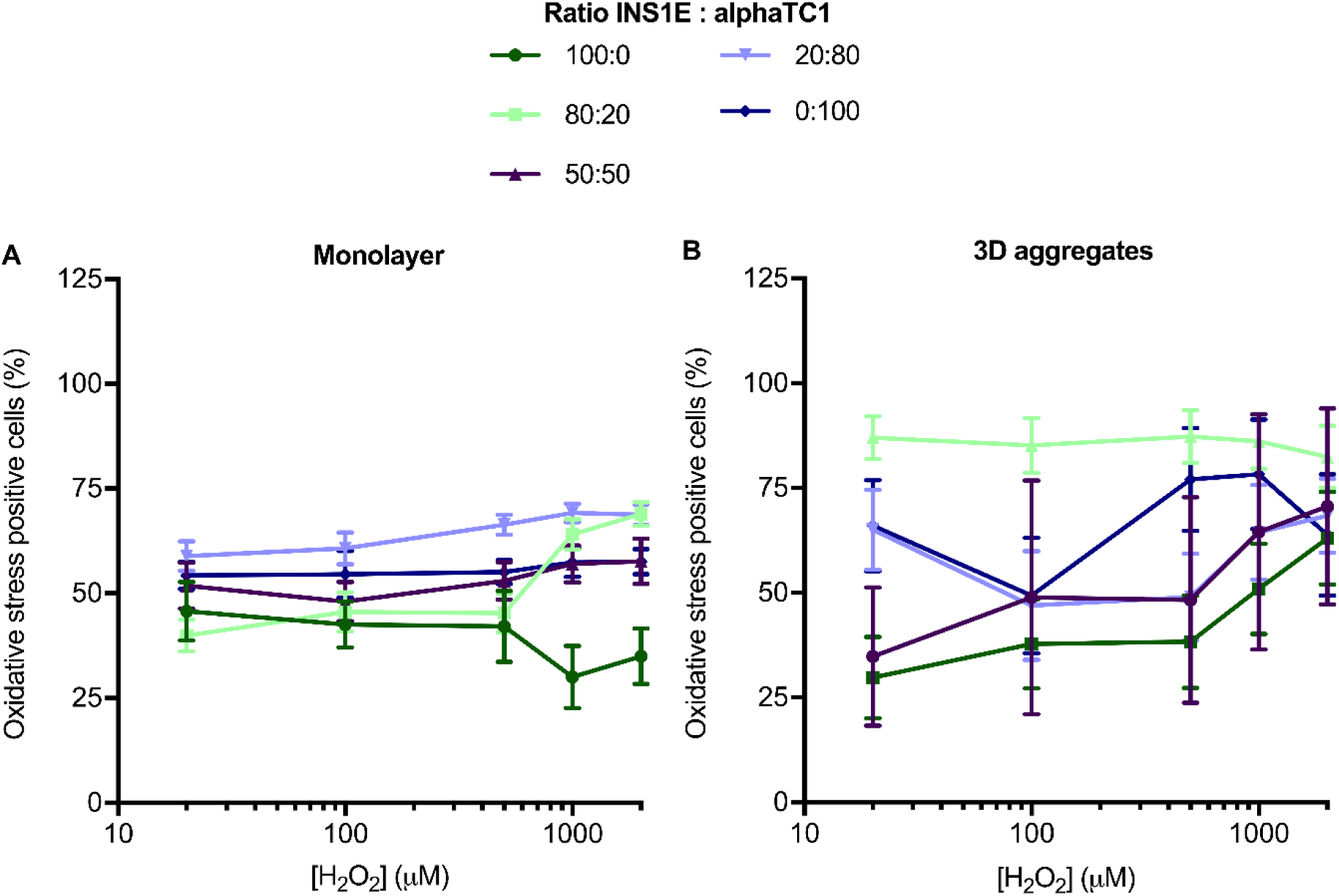
Total oxidative stress in monolayer culture and 3D aggregates. A) In a monolayer, oxidative stress induced by H_2_O_2_ increased the percentage of oxidative stress–positive cells in the ratios of 80:20 and 20:80 INS1E:alphaTC1. B) In 3D aggregates, oxidative stress induced by H_2_O_2_ did not increase the total percentage of oxidative stress–positive cells in the ratios 100:0, 80:20, 50:50, 20:80 and 0:100 INS1E:alphaTC1. N=3, A) n≥1; B) n≥2 and data are presented as mean ± SEM. All *p*-values are shown in Tables S3 and S4.

**Figure S4:**
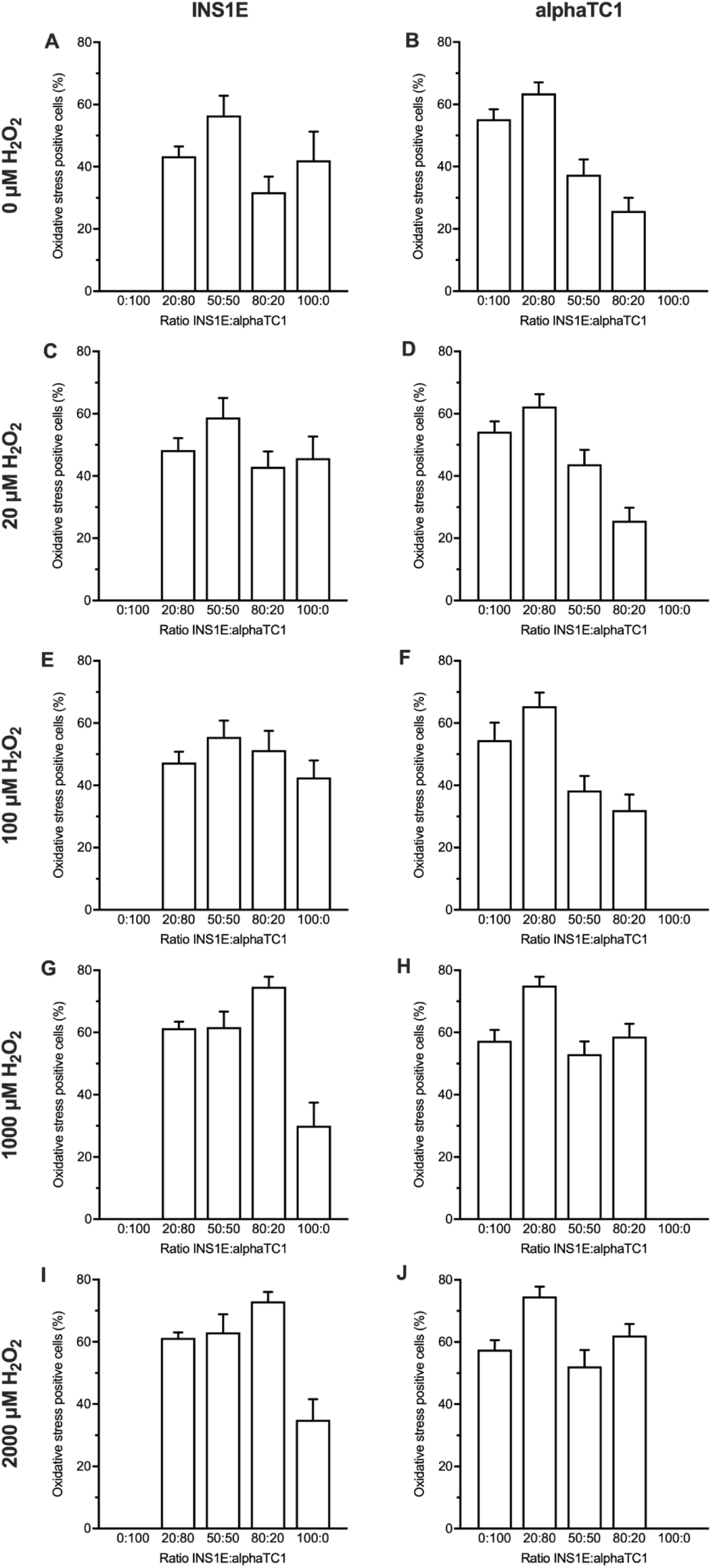
Oxidative stress–positive INS1E cells and alphaTC1 cells in monolayer culture when exposed to 0, 20, 100, 1000, and 2000 μM H_2_O_2_. *p*-values are shown in Tables S5 and S6.

**Figure S5:**
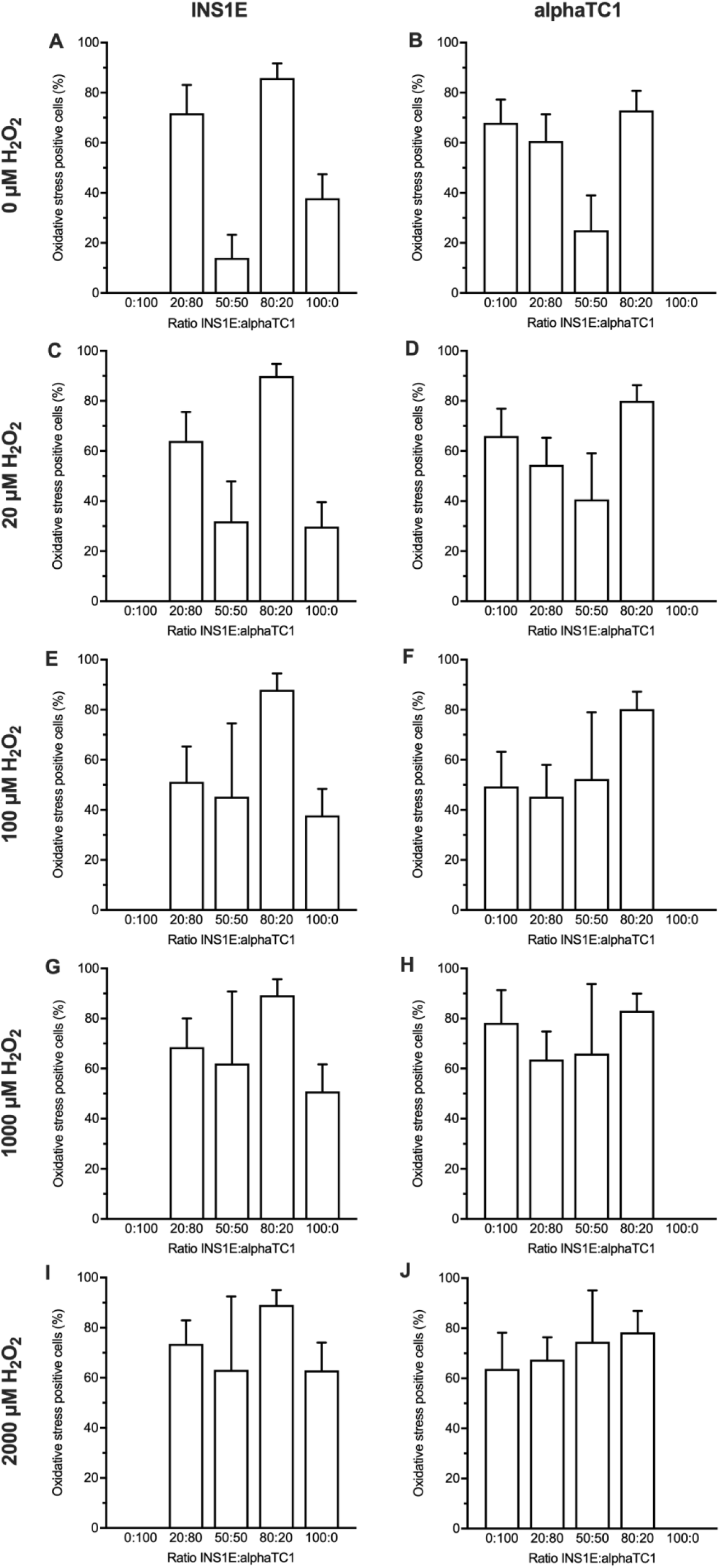
Oxidative stress–positive INS1E cells and alphaTC1 cells in 3D aggregates when exposed to 0, 20, 100, 1000 and 2000 μM H_2_O_2_. All *p*-values are shown in Tables S8 and S9.

## References

1. Barra JM, Tse HM. Redox-dependent inflammation in islet transplantation rejection. Front Endocrinol (Lausanne). 2018;9:175. Epub 2018/05/10. doi: 10.3389/fendo.2018.00175. PubMed PMID: 29740396; PubMed Central PMCID: PMCPMC5924790.

2. Saha SK, Lee SB, Won J, Choi HY, Kim K, Yang GM, et al. Correlation between oxidative stress, nutrition, and cancer initiation. Int J Mol Sci. 2017;18(7). Epub 2017/07/18. doi: 10.3390/ijms18071544. PubMed PMID: 28714931; PubMed Central PMCID: PMCPMC5536032.

3. Trachootham D, Lu W, Ogasawara MA, Nilsa RD, Huang P. Redox regulation of cell survival. Antioxid Redox Signal. 2008;10(8):1343–74. doi: 10.1089/ars.2007.1957. PubMed PMID: 18522489; PubMed Central PMCID: PMCPMC2932530.

4. Ursini F, Maiorino M, Forman HJ. Redox homeostasis: the golden mean of healthy living. Redox Biol. 2016;8:205–15. doi: 10.1016/j.redox.2016.01.010. PubMed PMID: 26820564; PubMed Central PMCID: PMCPMC4732014.

5. Sthijns MM, Randall MJ, Bast A, Haenen GR. Adaptation to acrolein through upregulating the protection by glutathione in human bronchial epithelial cells: the materialization of the hormesis concept. Biochem Biophys Res Commun. 2014;446(4):1029–34. doi: 10.1016/j.bbrc.2014.03.081. PubMed PMID: 24667599.

6. Sthijns MM, Thongkam W, Albrecht C, Hellack B, Bast A, Haenen GR, et al. Silver nanoparticles induce hormesis in A549 human epithelial cells. Toxicol In Vitro. 2017;40:223–33. doi: 10.1016/j.tiv.2017.01.010. PubMed PMID: 28109747.

7. Ahmed Alfar E, Kirova D, Konantz J, Birke S, Mansfeld J, Ninov N. Distinct levels of reactive oxygen species coordinate metabolic activity with beta-cell mass plasticity. Sci Rep. 2017;7(1):3994. doi: 10.1038/s41598-017-03873-9. PubMed PMID: 28652605; PubMed Central PMCID: PMCPMC5484671.

8. Pi J, Bai Y, Zhang Q, Wong V, Floering LM, Daniel K, et al. Reactive oxygen species as a signal in glucose-stimulated insulin secretion. Diabetes. 2007;56(7):1783–91. doi: 10.2337/db06-1601. PubMed PMID: 17400930.

9. Khacho M, Slack RS. Mitochondrial and reactive oxygen species signaling coordinate stem cell fate decisions and life long maintenance. Antioxid Redox Signal. 2018;28(11):1090–101. doi: 10.1089/ars.2017.7228. PubMed PMID: 28657337.

10. Khacho M, Clark A, Svoboda DS, Azzi J, MacLaurin JG, Meghaizel C, et al. Mitochondrial dynamics impacts stem cell identity and fate decisions by regulating a nuclear transcriptional program. Cell Stem Cell. 2016;19(2):232–47. Epub 2016/05/31. doi: 10.1016/j.stem.2016.04.015. PubMed PMID: 27237737.

11. Heinis M, Simon MT, Ilc K, Mazure NM, Pouyssegur J, Scharfmann R, et al. Oxygen tension regulates pancreatic beta-cell differentiation through hypoxia-inducible factor 1alpha. Diabetes. 2010;59(3):662–9. doi: 10.2337/db09-0891. PubMed PMID: 20009089; PubMed Central PMCID: PMCPMC2828660.

12. Cappelli APG, Zoppi CC, Silveira LR, Batista TM, Paula FM, da Silva PMR, et al. Reduced glucose-induced insulin secretion in low-protein-fed rats is associated with altered pancreatic islets redox status. J Cell Physiol. 2018;233(1):486–96. doi: 10.1002/jcp.25908. PubMed PMID: 28370189.

13. Robertson RP, Harmon J, Tran PO, Tanaka Y, Takahashi H. Glucose toxicity in beta-cells: type 2 diabetes, good radicals gone bad, and the glutathione connection. Diabetes. 2003;52(3):581–7. doi: 10.2337/diabetes.52.3.581. PubMed PMID: 12606496.

14. Ma Z, Moruzzi N, Catrina SB, Grill V, Bjorklund A. Hyperoxia inhibits glucose-induced insulin secretion and mitochondrial metabolism in rat pancreatic islets. Biochem Biophys Res Commun. 2014;443(1):223–8. doi: 10.1016/j.bbrc.2013.11.088. PubMed PMID: 24299957.

15. Maulucci G, Daniel B, Cohen O, Avrahami Y, Sasson S. Hormetic and regulatory effects of lipid peroxidation mediators in pancreatic beta cells. Mol Aspects Med. 2016;49:49–77. doi: 10.1016/j.mam.2016.03.001. PubMed PMID: 27012748.

16. Liu X, Han S, Yang Y, Kang J, Wu J. Glucose-induced glutathione reduction in mitochondria is involved in the first phase of pancreatic beta-cell insulin secretion. Biochem Biophys Res Commun. 2015;464(3):730–6. doi: 10.1016/j.bbrc.2015.07.016. PubMed PMID: 26164230.

17. Martin MA, Ramos S, Cordero-Herrero I, Bravo L, Goya L. Cocoa phenolic extract protects pancreatic beta cells against oxidative stress. Nutrients. 2013;5(8):2955–68. doi: 10.3390/nu5082955. PubMed PMID: 23912326; PubMed Central PMCID: PMCPMC3775237.

18. Liu CW, Bramer L, Webb-Robertson BJ, Waugh K, Rewers MJ, Zhang Q. Temporal expression profiling of plasma proteins reveals oxidative stress in early stages of type 1 diabetes progression. J Proteomics. 2018;172:100–10. doi: 10.1016/j.jprot.2017.10.004. PubMed PMID: 28993202; PubMed Central PMCID: PMCPMC5726913.

19. Miki A, Ricordi C, Sakuma Y, Yamamoto T, Misawa R, Mita A, et al. Divergent antioxidant capacity of human islet cell subsets: a potential cause of beta-cell vulnerability in diabetes and islet transplantation. PLoS One. 2018;13(5):e0196570. doi: 10.1371/journal.pone.0196570. PubMed PMID: 29723228; PubMed Central PMCID: PMCPMC5933778.

20. Stancill JS, Broniowska KA, Oleson BJ, Naatz A, Corbett JA. Pancreatic beta-cells detoxify H_2_O_2_ through the peroxiredoxin/thioredoxin antioxidant system. J Biol Chem. 2019;294(13):4843–53. doi: 10.1074/jbc.RA118.006219. PubMed PMID: 30659092; PubMed Central PMCID: PMCPMC6442057.

21. Mehmeti I, Lortz S, Avezov E, Jorns A, Lenzen S. ER-resident antioxidative GPx7 and GPx8 enzyme isoforms protect insulin-secreting INS-1E beta-cells against lipotoxicity by improving the ER antioxidative capacity. Free Radic Biol Med. 2017;112:121–30. doi: 10.1016/j.freeradbiomed.2017.07.021. PubMed PMID: 28751022.

22. Lortz S, Gurgul-Convey E, Naujok O, Lenzen S. Overexpression of the antioxidant enzyme catalase does not interfere with the glucose responsiveness of insulin-secreting INS-1E cells and rat islets. Diabetologia. 2013;56(4):774–82. doi: 10.1007/s00125-012-2823-7. PubMed PMID: 23306382.

23. Mehmeti I, Lortz S, Elsner M, Lenzen S. Peroxiredoxin 4 improves insulin biosynthesis and glucose-induced insulin secretion in insulin-secreting INS-1E cells. J Biol Chem. 2014;289(39):26904–13. doi: 10.1074/jbc.M114.568329. PubMed PMID: 25122762; PubMed Central PMCID: PMCPMC4175331.

24. Sneddon JB, Tang Q, Stock P, Bluestone JA, Roy S, Desai T, et al. Stem cell therapies for treating diabetes: progress and remaining challenges. Cell Stem Cell. 2018;22(6):810–23. Epub 2018/06/03. doi: 10.1016/j.stem.2018.05.016. PubMed PMID: 29859172; PubMed Central PMCID: PMCPMC6007036.

25. Lebreton F, Lavallard V, Bellofatto K, Bonnet R, Wassmer CH, Perez L, et al. Insulin-producing organoids engineered from islet and amniotic epithelial cells to treat diabetes. Nat Commun. 2019;10(1):4491. Epub 2019/10/05. doi: 10.1038/s41467-019-12472-3. PubMed PMID: 31582751; PubMed Central PMCID: PMCPMC6776618.

26. Chen L, Wu M, Jiang S, Zhang Y, Li R, Lu Y, et al. Skin toxicity assessment of silver nanoparticles in a 3D epidermal model compared to 2D keratinocytes. Int J Nanomedicine. 2019;14:9707–19. doi: 10.2147/IJN.S225451. PubMed PMID: 31849463; PubMed Central PMCID: PMCPMC6910103.

27. Lehmann R, Zuellig RA, Kugelmeier P, Baenninger PB, Moritz W, Perren A, et al. Superiority of small islets in human islet transplantation. Diabetes. 2007;56(3):594–603. doi: 10.2337/db06-0779. PubMed PMID: 17327426.

28. Rahman I, Kode A, Biswas SK. Assay for quantitative determination of glutathione and glutathione disulfide levels using enzymatic recycling method. Nat Protoc. 2006;1(6):3159–65. doi: 10.1038/nprot.2006.378. PubMed PMID: 17406579.

29. Deglasse JP, Roma LP, Pastor-Flores D, Gilon P, Dick TP, Jonas JC. Glucose acutely reduces cytosolic and mitochondrial H_2_O_2_ in rat pancreatic beta cells. Antioxid Redox Signal. 2019;30(3):297–313. doi: 10.1089/ars.2017.7287. PubMed PMID: 29756464.

30. Myasnikova D, Osaki T, Onishi K, Kageyama T, Zhang Molino B, Fukuda J. Synergic effects of oxygen supply and antioxidants on pancreatic beta-cell spheroids. Sci Rep. 2019;9(1):1802. doi: 10.1038/s41598-018-38011-6. PubMed PMID: 30755634; PubMed Central PMCID: PMCPMC6372787.

31. Komatsu H, Cook C, Wang CH, Medrano L, Lin H, Kandeel F, et al. Oxygen environment and islet size are the primary limiting factors of isolated pancreatic islet survival. PLoS One. 2017;12(8):e0183780. doi: 10.1371/journal.pone.0183780. PubMed PMID: 28832685; PubMed Central PMCID: PMCPMC5568442.

32. Gerber PA, Rutter GA. The role of oxidative stress and hypoxia in pancreatic beta-cell dysfunction in diabetes mellitus. Antioxid Redox Signal. 2017;26(10):501–18. doi: 10.1089/ars.2016.6755. PubMed PMID: 27225690; PubMed Central PMCID: PMCPMC5372767.

33. Joshi S, Kumar S, Ponnusamy MP, Batra SK. Hypoxia-induced oxidative stress promotes MUC4 degradation via autophagy to enhance pancreatic cancer cells survival. Oncogene. 2016;35(45):5882–92. doi: 10.1038/onc.2016.119. PubMed PMID: 27109098; PubMed Central PMCID: PMCPMC5079846.

34. Sthijns M, van Blitterswijk CA, LaPointe VLS. Redox regulation in regenerative medicine and tissue engineering: the paradox of oxygen. J Tissue Eng Regen Med. 2018;12(10):2013–20. doi: 10.1002/term.2730. PubMed PMID: 30044552; PubMed Central PMCID: PMCPMC6221092.

35. Nagar R, Khan AR, Poonia A, Mishra PK, Singh S. Metabolism of cisplatin in the organs of Rattus norvegicus: role of Glutathione S-transferase P1. Eur J Drug Metab Pharmacokinet. 2015;40(1):45–51. doi: 10.1007/s13318-014-0176-y. PubMed PMID: 24474500.

36. Padmasekar M, Lingwal N, Samikannu B, Chen C, Sauer H, Linn T. Exendin-4 protects hypoxic islets from oxidative stress and improves islet transplantation outcome. Endocrinology. 2013;154(4).

37. Rodriguez-Diaz R, Molano RD, Weitz JR, Abdulreda MH, Berman DM, Leibiger B, et al. Paracrine interactions within the pancreatic islet determine the glycemic set point. Cell Metab. 2018;27(3):549–58 e4. doi: 10.1016/j.cmet.2018.01.015. PubMed PMID: 29514065; PubMed Central PMCID: PMCPMC5872154.

38. Chandravanshi B, Bhonde RR. Shielding engineered islets with mesenchymal stem cells enhance survival under hypoxia. J Cell Biochem. 2017;118(9):2672–83. doi: 10.1002/jcb.25885. PubMed PMID: 28098405.

39. Li X, Lang H, Li B, Zhang C, Sun N, Lin J, et al. Change in viability and function of pancreatic islets after coculture with mesenchymal stromal cells: a systemic review and meta-analysis. J Diabetes Res. 2020;2020:5860417. doi: 10.1155/2020/5860417. PubMed PMID: 32309447; PubMed Central PMCID: PMCPMC7132593.

40. Laporte C, Tubbs E, Cristante J, Gauchez AS, Pesenti S, Lamarche F, et al. Human mesenchymal stem cells improve rat islet functionality under cytokine stress with combined upregulation of heme oxygenase-1 and ferritin. Stem Cell Res Ther. 2019;10(1):85. doi: 10.1186/s13287-019-1190-4. PubMed PMID: 30867050; PubMed Central PMCID: PMCPMC6416979.

41. Lammert E, Thorn P. The role of the islet niche on beta cell structure and function. J Mol Biol. 2020;432(5):1407–18. doi: 10.1016/j.jmb.2019.10.032. PubMed PMID: 31711959.

42. Allagnat F, Klee P, Cardozo AK, Meda P, Haefliger JA. Connexin36 contributes to INS-1E cells survival through modulation of cytokine-induced oxidative stress, ER stress and AMPK activity. Cell Death Differ. 2013;20(12):1742–52. doi: 10.1038/cdd.2013.134. PubMed PMID: 24096873; PubMed Central PMCID: PMCPMC3824597.

43. McCall M, Shapiro AM. Update on islet transplantation. Cold Spring Harb Perspect Med. 2012;2(7):a007823. doi: 10.1101/cshperspect.a007823. PubMed PMID: 22762022; PubMed Central PMCID: PMCPMC3385934.

44. Shapiro AM, Ricordi C, Hering BJ, Auchincloss H, Lindblad R, Robertson RP, et al. International trial of the Edmonton protocol for islet transplantation. N Engl J Med. 2006;355(13):1318–30. doi: 10.1056/NEJMoa061267. PubMed PMID: 17005949.

45. Jiang J, Wang K, Nice EC, Zhang T, Huang C. High-throughput screening of cellular redox sensors using modern redox proteomics approaches. Expert Rev Proteomics. 2015;12(5):543–55. doi: 10.1586/14789450.2015.1069189. PubMed PMID: 26184698.

46. Sthijns M, Jetten MJ, Mohammed SG, Claessen SMH, de Vries RHW, Stell A, et al. Oxidative stress in pancreatic alpha and beta cells as a selection criterion for biocompatible biomaterials. Biomaterials. 2021;267:120449. Epub 2020/11/01. doi: 10.1016/j.biomaterials.2020.120449. PubMed PMID: 33129188.

47. Veres A, Faust AL, Bushnell HL, Engquist EN, Kenty JH, Harb G, et al. Charting cellular identity during human in vitro beta-cell differentiation. Nature. 2019;569(7756):368–73. doi: 10.1038/s41586-019-1168-5. PubMed PMID: 31068696; PubMed Central PMCID: PMCPMC6903417.

